# Multipolar spindle assembly and mitotic slippage underlie symbiont-mediated asexual reproduction

**DOI:** 10.64898/2026.06.11.731681

**Authors:** Laura C. Fricke, Michael A. Gelaw, Jack A. Culotta, Amelia R. I. Lindsey

**Affiliations:** Department of Entomology, University of Minnesota, St. Paul, Minnesota, 55108

**Keywords:** microtubule organizing center, MTOC, centrosome, parthenogenesis, host-microbe interaction, symbiosis, *Wolbachia*

## Abstract

Maternally inherited bacteria can profoundly impact animal biology. A notable example is the induction of thelytokous parthenogenesis: the asexual production of female offspring. Despite decades of symbiont-mediated parthenogenesis documented across diverse arthropods, the underlying cell biological mechanisms remain poorly understood. In some species of parasitic wasps, *Wolbachia* causes a chromosome segregation failure during the first mitotic division in unfertilized haploid embryos. This diplodizes the embryo and facilitates asexual reproduction of females. To elucidate the mechanism of mitotic failure and understand how the animal continues development, we compared meiosis and early embryonic mitoses of the parasitoid wasp, *Trichogramma pretiosum*, with and without their parthenogenesis-inducing *Wolbachia* symbiont. We show that meiosis in these animals is anastral, and embryos undergo *de novo* microtubule organizing center (MTOC) biogenesis prior to the first embryonic mitosis, regardless of *Wolbachia*. During the first mitosis, embryos with *Wolbachia* had increased numbers of MTOCs associated with the oocyte nucleus and formed multipolar spindles in a modified metaphase, after which chromosome segregation failed. A restitution nucleus formed, and mitotic slippage enabled the diploid nucleus to transition to interphase and replicate. Subsequent mitotic divisions inherited supernumerary MTOCs, but formed pseudo-bipolar spindles and segregated normally. These results demonstrate that *Wolbachia*-mediated parthenogenesis stems from altered MTOC biology, provide a structural basis for mitotic failure and the subsequent restitution. Symbiont-mediated parthenogenesis offers unique opportunities to uncover the cell biology of asexual reproduction and a fascinating model for understanding mitosis more broadly.

## INTRODUCTION

The accurate replication and segregation of genetic material is essential for all life. Disruptions to the timing of the cell cycle or dynamics of chromosome segregation can lead to numerous defects including aneuploidy, genomic instability, mitotic catastrophe, and cell death (1). Such defects are characteristic of cancer cells and can also be induced by pathogens such as *Chlamydia, Listeria monocytogenes,* and myriad viruses (2–8). In contrast, some strains of the bacterium *Wolbachia* (Alphaproteobacteria: Rickettsiales) have evolved to cause a mitotic failure in certain insect embryos that is not catastrophic, but instead facilitates thelytokous parthenogenesis (the asexual reproduction of female offspring) a phenotype known as parthenogenesis-induction (PI)(9, 10).

The maternally transmitted intracellular bacterium *Wolbachia* infects approximately half of all insect species (11), and are most well-known for host reproductive manipulations that confer advantages for *Wolbachia-*infected, and thus *Wolbachia*-transmitting, females in the population (12, 13). For example, male-killing, conditional sperm-egg incompatibilities, feminization, and the above-mentioned PI, maximize transmission of *Wolbachia* through generations (12, 13). Importantly, PI occurs in animals such as parasitic wasps like *Trichogramma* (Fig. 1A) in which sex is determined by ploidy (i.e. haplodiploidy) (14, 15). In the absence of *Wolbachia,* females are produced from fertilized (i.e., diploid) embryos, and males from unfertilized (i.e., haploid) embryos (Fig. 1B). Under one form of PI, *Wolbachia* targets the first zygotic mitosis of unfertilized embryos, effectively preventing sister chromatid segregation, resulting in diploidization (10, 16). Thus, *Wolbachia* leverages the host sex determination system in favor of its own propagation, diplodizing the unfertilized, male-destined embryo to instead facilitate female development and propagation of *Wolbachia* (Fig. 1B) (10). Remarkably, in these embryos subsequent mitotic cycles are successful, despite the initial failure. Chromosome staining of early-stage wasp embryos revealed that embryonic mitosis fails only in the first nuclear cycle (NC) (10, 16). Mitosis is initiated, but the chromosomes do not segregate during anaphase of NC1, and a diploid “restitution” nucleus is formed (Fig. 1C). The diploid restitution nucleus then enters interphase, chromosomes replicate, and the now 4n nucleus successfully divides into two 2n daughter nuclei.

**Figure 1.**
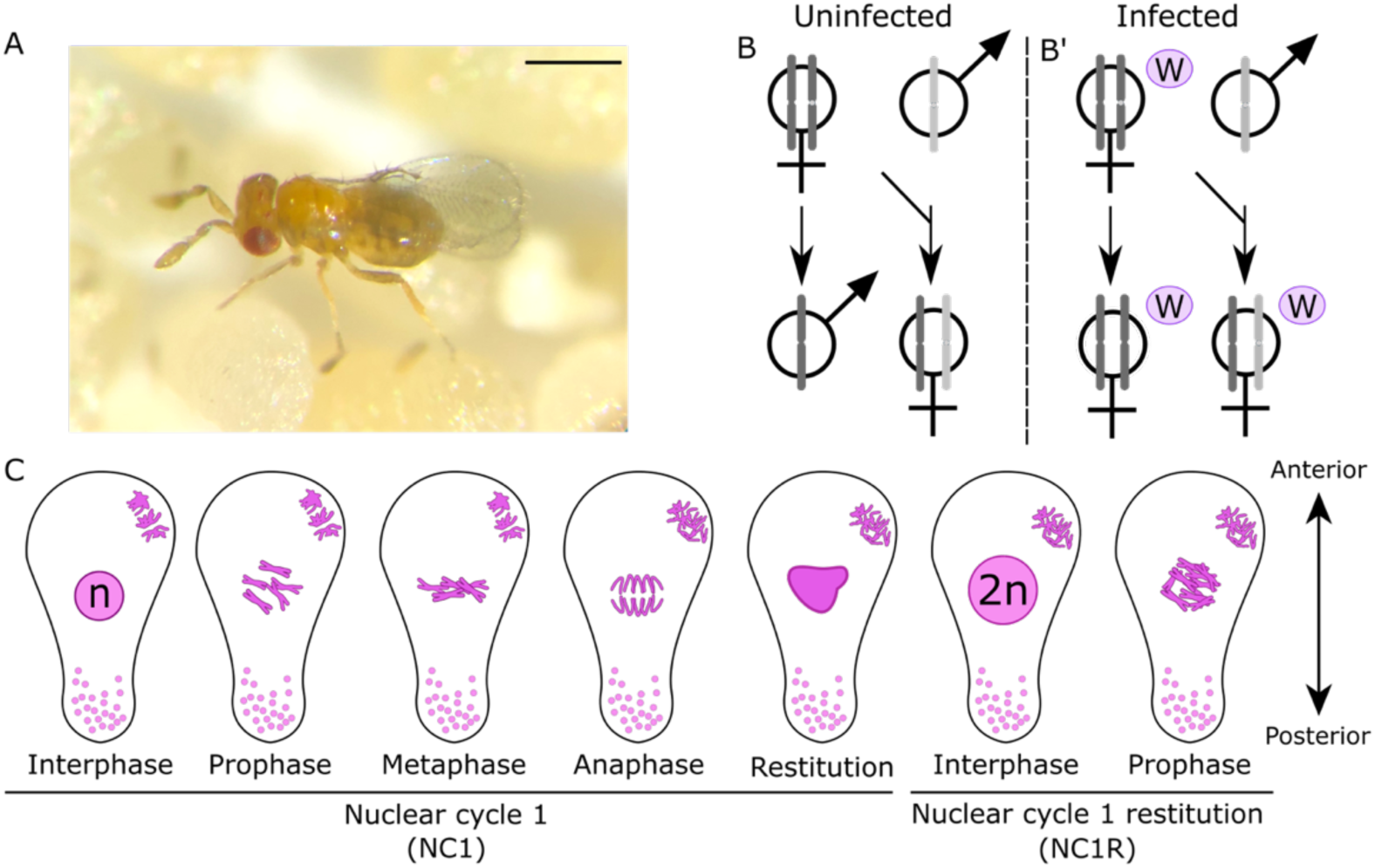
*Wolbachia* induces gamete duplication in embryos of the parasitic wasp, *Trichogramma.* **(A)** *Trichogramma pretiosum* adult female, parasitizing *Ephestia kuehniella* eggs. The scale bar represents 150 μm. **(B)** Schematic showing haplodiploid sex determination in the absence of *Wolbachia*, where males develop from haploid eggs, and females from fertilized, diploid eggs. **(B’)** Model of haplodiploid sex determination in the presence of parthenogenesis inducing *Wolbachia*. In *Trichogramma*, unfertilized eggs are diploidized by *Wolbachia,* resulting in homozygous female offspring, while fertilized eggs yield diploid, heterozygous females. Both types of female offspring inherit *Wolbachia*. **(C)** Schematic of the diploidization process proposed by Stouthamer and Kazmer (1994) (10). Unfertilized embryos complete meiosis and produce 3 polar bodies at the cortex and one haploid pronucleus of five chromosomes at the center. The interphase pronucleus replicates and enters prophase of mitosis I with 5 condensed pairs of chromosomes (n=10). In metaphase, chromosomes align classically, however sister chromatids fail to separate during anaphase, resulting in a diploid restitution nucleus. The restitution nucleus enters interphase in which DNA is replicated (to n=20) and the first diploid restitution mitosis, (1R) begins with 10 pairs of chromosomes. Puncta at the posterior of embryos represent *Wolbachia*.

While diploidization due to *Wolbachia* is clear, the mechanisms responsible have remained unknown since this discovery was made more than 30 years ago. *Wolbachia*-mediated mitotic failure has a strict set of conditions that make this a valuable study system for understanding the intricacies of mitosis. For example, for PI to be successful, *Wolbachia* cannot cause indiscriminate diploidization: a specific mitosis needs to be targeted. Furthermore, the impacts on this specific mitosis cannot persist; a challenge given that many mitotic defects are imprinted into subsequent cycles (1). Finally, because symbiont-mediated PI is not necessarily ubiquitous within a species or population, matings between PI-*Wolbachia* infected females and *Wolbachia*-uninfected males are relatively common in some areas (10, 15, 17). In such cases, *Wolbachia* does not impact the first mitosis if the embryo is fertilized (10, 17). Thus, the mechanism of mitotic failure must be fertilization-sensitive. In contrast to the pathogens that evolved mechanisms for “breaking” host mitosis (3, 5–8), PI-*Wolbachia* evolved tools to modulate a sensitive process and avoid irreversible damage to the host.

In this study, we developed protocols for immunofluorescence staining of embryos of *Trichogramma*, parasitic wasps where PI-*Wolbachia* were first described (9), to examine spindle and chromosome dynamics. We then characterized meiosis and early embryonic mitoses of wasp embryos with and without *Wolbachia* infections. In the first zygotic mitosis of *Wolbachia*-infected embryos, chromatid segregation failure is associated with a disruption of microtubule organizing center (MTOC) dynamics and the development of a multipolar spindle. The mitosis fails, and mitotic slippage (exiting mitosis without undergoing chromosome segregation) enables the nucleus to enter interphase, ultimately resulting in diploidization. These discoveries offer insight into a fascinating instance of conditional mitotic failure and the mechanisms of asexual reproduction.

## METHODS

### Insect lines

*Trichogramma pretiosum* were maintained at 25°C on a 24-h, 12:12 light:dark cycle. Wasps were hosted on UV-irradiated eggs of *Ephestia kuehniella* (Koppert Global or Bioline Agrosciences depending on availability). *Ephestia kuehniella* eggs were provided to wasps on strips of double-sided tape adhered to cardstock (i.e., egg cards). To control for differences in genetic background, we leveraged a previously generated pair of colonies that only differ in their *Wolbachia* infection status (18). These colonies are derived from the naturally *Wolbachia* infected, asexual “Insectary” colony, which originated from a single *Wolbachia*-infected female collected in Peru (19, 20), and the naturally *Wolbachia-*uninfected, sexually reproducing “CA29” colony, derived from a single *Wolbachia*-uninfected son-mated female collected in California, USA (21). Because many *Wolbachia*-infected *T. pretiosum* have undergone a significant loss of sexual function due to mutations in the nuclear genome (15, 20), they cannot be cured with antibiotics and propagated without their *Wolbachia* infection (22). To overcome this, genetically matched colonies were generated by introgressing the CA29 nuclear genome (that has complete sexual function) into the Insectary cytoplasm via seven generations of backcrossing to produce the *Wolbachia*-infected colony “Introgressed isofemale Line B” (IILB+) (18). IILB+ was then split in two and the complementary *Wolbachia*-uninfected line (IILB-) was generated via antibiotic treatment (18). Both colonies have been maintained in parallel since 2016. Genome sequencing indicated that the nuclear genomes of these two introgressed lines are identical and >99% CA29 in origin (18). *Drosophila melanogaster* samples were used as positive controls in western blots (see below), and leveraged flies from a *Wolbachia*-free version of the stock DGRP-352 (RRID:BDSC_83728) produced and maintained following published approaches (23).

### Validation of reproductive mode and *Wolbachia* infection

The reproductive mode and *Wolbachia* infection status of colonies was validated using standard methods (24) prior to all experiments. To test for the asexual reproduction of females, parasitized *Ephestia kuehniella* eggs (each containing a single late-stage *Trichogramma* pupa) were removed from egg cards with a dampened fine tipped paint brush and isolated into separate 12x75 mm glass rearing vials (Avantor) plugged with cotton. Once unmated females emerged, they were given a small egg card, and the production of female offspring was used to verify thelytokous parthenogenesis, consistent with *Wolbachia*-mediated PI (24). To assess *Wolbachia* infection status, colonies were periodically screened for *Wolbachia* DNA following standard methods (24). DNA extractions were performed on individual adult wasps using a HotShot method (25) adapted for *Trichogramma*, in a total reaction volume of 24 µL (26). PCRs were performed using Q5 Hot Start High-Fidelity 2x Master Mix (New England Biolabs) with 500 nM of each *Wolbachia* specific primer (W-specF and W-specR, (27)) and 1 µL of template DNA per reaction. Products were run on a 1% agarose gel alongside a 100 base pair ladder (New England Biolabs). The gel was stained with 3X GelRed (Biotium) in deionized water and visualized under UV light.

### Collection of unfertilized embryos

To collect *Trichogramma* embryos, newly emerged (less than 24h) adult females were provided *ad libitum Ephestia kuehniella* eggs and allowed to parasitize for a defined time window (see results). To harvest *Wolbachia*-uninfected, haploid embryos, females from the *Wolbachia*-uninfected colony IILB- were isolated as pupae to prevent mating and pooled upon emergence prior to hosting on *Ephestia kuehniella*.

### Embryo fixation and immunofluorescence

Parasitized *Ephestia kuehniella* eggs were lysed in cold 1X phosphate buffered saline (PBS) with a disposable pestle in 1.7 mL microcentrifuge tubes to release *Trichogramma* embryos. To remove large host debris, lysate was filtered through a 70 µm cell strainer (GenClone). Embryos in the filtrate were moved to fresh 1.7 mL microcentrifuge tubes and centrifuged for 15 seconds at 5,000 rpm. Excess liquid was removed without disturbing the pelleted embryos, after which they were resuspended in 200 µL of fixative solution comprised of 4% paraformaldehyde and 1% NP40 in 1X PBS. 400 µL of heptane was added and embryos were nutated for 30 minutes at room temperature. Heptane was discarded, embryos were pelleted by centrifugation for 15 seconds at 5,000 rpm, and the fixative was removed. Embryos were resuspended in 400 µL of methanol and 500 µL of heptane. To permeabilize the chorion and vitelline membrane, samples were “cracked” by repeatedly striking the microcentrifuge tube against the laboratory bench (approximately twice every second) for 2 minutes. Samples were allowed to rest for 1 minute then subjected to a second round of cracking. Heptane was removed and the embryos were sonicated for 3 seconds at level 5 with the XL-2000 Misonix Sonicator fit with the standard probe. The sonicated embryos were transferred to glass spot plates and washed three times with 200 µL of methanol, followed by three more washes with 200 µL PBS with 1% Tween 20 (PBST). Embryos were then incubated overnight at 4°C in 250 µL PBST with 5% bovine serum albumin (PBST-BSA). The PBST-BSA was removed and replaced with fresh PBST-BSA containing primary antibodies against α-tubulin and γ-tubulin at a1:200 dilution (Supplemental Table S1), then incubated overnight at 4°C. Embryos were washed three times with PBST, resuspended in secondary antibodies (Supplemental Table S1) diluted in PBST-BSA at 1:1,000, then incubated for two hours at room temperature. Secondary antibodies were removed, and embryos were washed three times with PBST. Samples were mounted in 25 µL ProLong Glass Antifade Mountant with NucBlue Stain (18, 19) (Invitrogen) on slides bordered by one layer of standard scotch tape as a spacer.

### Microscopy and image analysis

Embryos were imaged on a Nikon AX R confocal microscope with a 60X oil immersion objective (N.A.=1.42). The galvanic scanner was used with lasers and detection filters for the 405, 488, and 561 nm excitation lines. Three-dimensional images were obtained using the Z-stack function with a step size of 0.2 µm. The number of steps varied depending on the size of the embryo and its positioning in the mounting medium. All images were processed with DenoiseAi and 3D Deconvolution in NIS Elements implemented via GA3 batch processing. Maximum intensity projections were generated for all figures. Mitotic staging was conducted manually using chromosome morphology, polar body formation, and cytoplasmic aster characteristics a guides (10, 28). Distances between γ-tubulin foci were measured with NIS-Elements 3D Object Measurement, with centroids defined as the focal point. Nuclear associated MTOCs were manually counted from the three-dimensional views, with α-tubulin signal used to verify connection with the nucleus.

### Western blots

Insect samples were lysed with a motorized pestle in 100 µL of 150 mM NaCl, 1% Triton X-100, and 50 mM TrisHCl (pH 8.0) containing 1% HALT protease inhibitor cocktail (Thermo Scientific). Samples were centrifuged for 15 seconds at 5,000 rpm, and 20 µL of the supernatant was combined 1:1 with 5% β-mercaptoethanol in 2X Laemmli sample buffer. The mixture was incubated for 5 minutes at 95°C. 35 µL of each sample and 10 µL of PageRuler Prestained Protein ladder (Thermo Scientific) was run on a 4-20% gradient polyacrylamide gel (BioRad) for 60 minutes at 140 V. The gel was transferred to a PVDF membrane using a Trans-blot Turbo transfer pack and Trans-blot Turbo cassette (BioRad) on the Turbo setting. The membrane was submerged in 10 mL of SuperBlock blocking buffer (blotting, in TBS) (Thermo Scientific) and incubated overnight at 4°C on an orbital shaker at 40 rpm. After removing the blocking buffer, 15 mL of primary antibody (Supplemental Table S1) diluted 1:1,000 in SuperBlock blocking buffer blotting in TBS + 0.05% Tween-20 was added to the membrane and nutated for 1 hour at room temperature. The primary antibody solution was removed, and the membrane was washed five times with 1X TBST (0.1% Tween-20) for ten minutes each time on an orbital shaker.

Membranes were incubated in HRP-conjugated secondary antibody (Supplemental Table S1) diluted 1:10,000 in 15 mL SuperBlock blocking buffer (blotting, in TBS) for 1 hour at room temperature on an orbital shaker. The secondary antibody was removed, and the membrane was washed five times in 1X TBST (0.1% Tween-20) for ten minutes each time. For antibody detection, a 1:1 ratio of Luminol/Enhancer to Stable Peroxide Buffer (Thermo Scientific) was added to the membrane and gently nutated by hand for 3 minutes. The membrane was imaged on a C-DiGit Blot scanner using the high sensitivity setting.

### Statistics and data visualization

Statistical analysis and data visualization were performed in R version 4.5.3 (R Core Team 2014). Significant differences in γ-tubulin intraplanar distance were assessed with a Mann-Whitney U test, and *p*-values were adjusted using the Bonferroni-Holm correction, with significance defined as *p_adj_* <0.05. Significant differences in the numbers of both MTOCs and γ-tubulin foci associated with the spindle were assessed with a Generalized Linear Model (GLM) that included unique combinations of *Wolbachia* status and NC as fixed effects, and a Poisson error distribution with a log link function. Schematics were created in Inkscape, and data visualization applied the “discrete rainbow colour scheme” designed by Paul Tol (https://personal.sron.nl/~pault/).

## RESULTS

### Development of immunofluorescence protocols for *Trichogramma* embryos

Prior to developing immunofluorescence staining protocols for *Trichogramma* embryos, we validated commercially available primary antibodies via western blotting. Primary antibodies against α- and γ-tubulin recognized *Trichogramma* proteins of the expected size (Supplemental Figures S1, S2). We then developed protocols for immunofluorescence in *Trichogramma* embryos to facilitate visualization of spindle and MTOC dynamics during meiosis and mitosis (Fig. 2,3). Counterstaining with NucBlue (Hoechst 33342) effectively captured chromosome morphology and highlighted individual *Wolbachia* cells as ∼0.7 µm puncta in the *Wolbachia-*infected embryos (Figs. 2,3).

**Figure 2.**
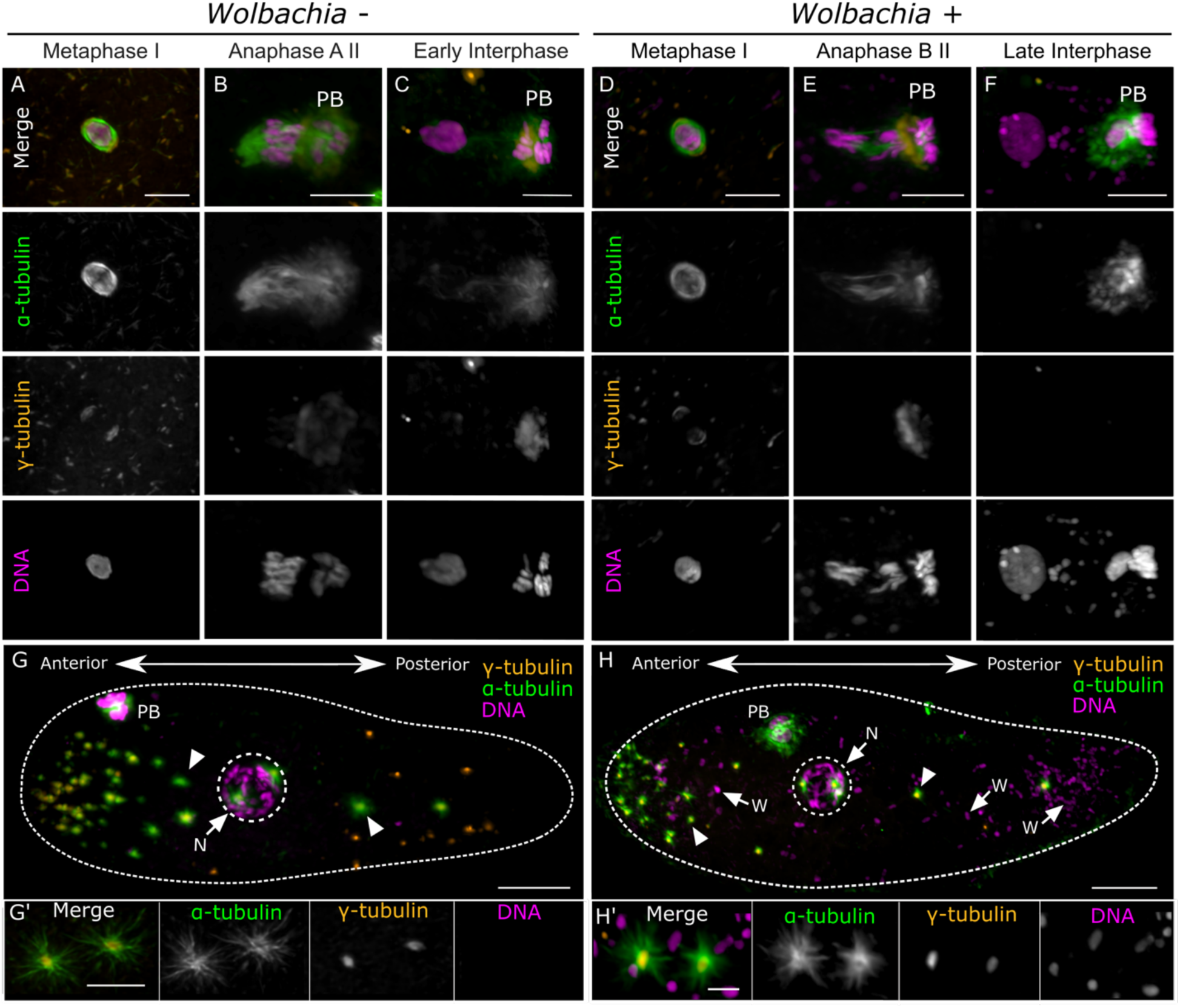
Meiosis in *Trichogramma* embryos is anastral and *de novo* MTOCs form during meiosis II. **(A-F)** 0-to 105-minute-old embryos were collected, fixed, and stained with NucBlue (Hoechst 33342) for DNA, and antibodies against α- and γ-tubulin. For each panel (a column of images), individual channels are shown in grey scale alongside the colorized composite. Scale bars represent 5 μm. **(A)** *Wolbachia*-uninfected embryo in metaphase of meiosis I **(B)** *Wolbachia*-uninfected embryo in anaphase A of meiosis II. Polar body 1 is labeled with PB. **(C)** *Wolbachia*-uninfected embryo in early interphase **(D)** *Wolbachia*-infected embryo in metaphase of meiosis I. **(E)** *Wolbachia*-infected embryo in anaphase B of meiosis II. Polar bodies are labeled with PB. **(F)** *Wolbachia*-infected embryo in interphase after the completion of meiosis. The pronucleus at the top is associated with *Wolbachia* foci. This image is a later time point than is shown in (C). **(G,H)** 0- to 120-minute-old embryos were collected, fixed, and stained with NucBlue (Hoechst 33342) for DNA, and antibodies against α- and γ-tubulin. Nuclei are denoted with dotted lines and the ‘N’ labeled arrow, polar bodies are denoted by ‘PB’, and select cytoplasmic asters are indicated with arrowheads. **(G)** *Wolbachia*-uninfected, unfertilized embryo in prophase of mitosis I. (G’) Cytoplasmic asters of α-tubulin contain central γ-tubulin foci. **(H)** A *Wolbachia-*infected embryo in prophase of mitosis I. Cytoplasmic puncta of DNA are *Wolbachia,* some of which are indicated with the ‘W’ labeled arrows. (H’) Cytoplasmic asters of α-tubulin contain central γ-tubulin foci. G and H are colorized composite images with scale bars representing 10 μm. Images in G’ and H’ show individual channels in grey scale in addition to the colorized composite with a scale bar of 5 μm.

**Figure 3.**
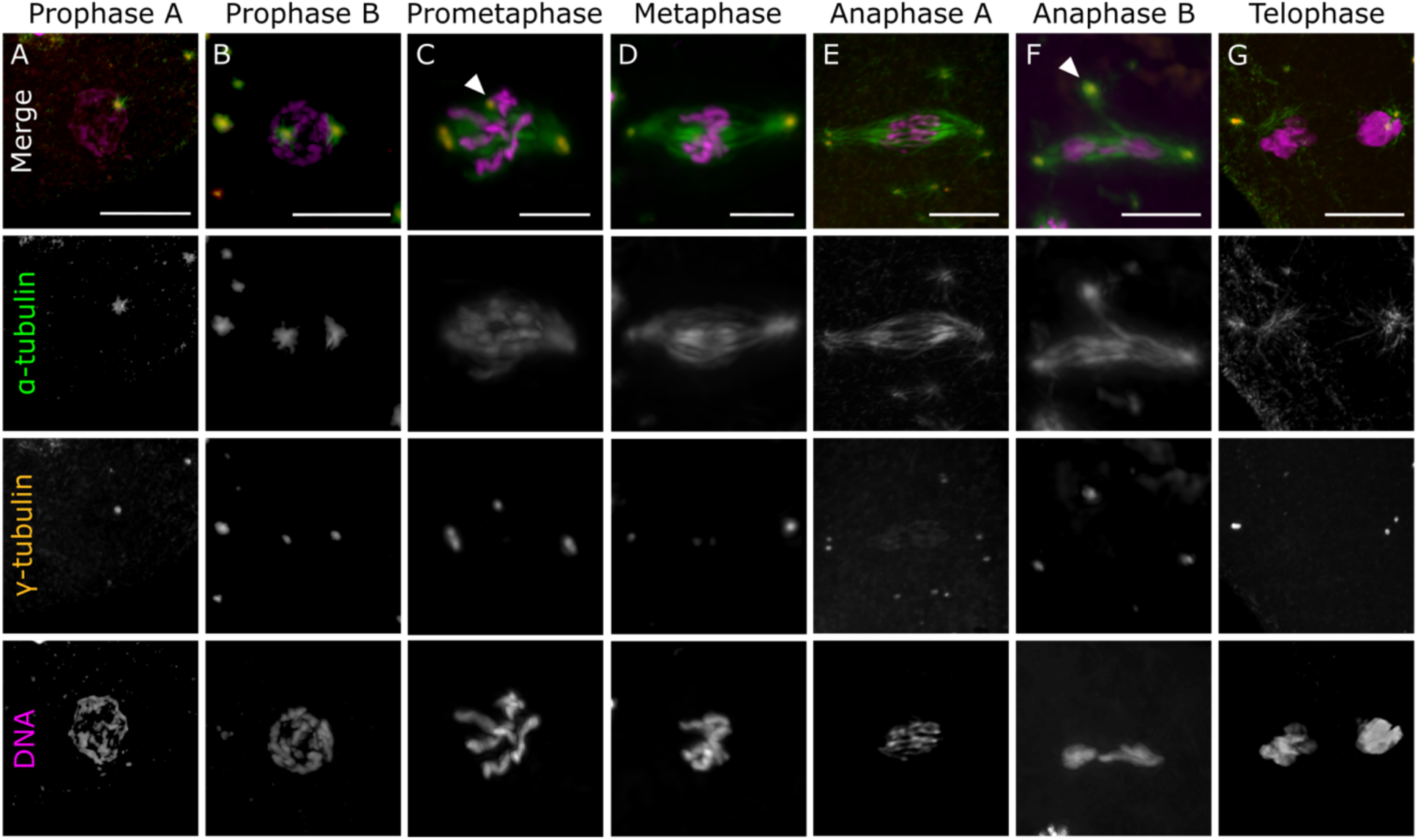
The first zygotic mitosis in *Wolbachia*-uninfected *Trichogramma* embryos is bipolar. 30-90-minute-old *Wolbachia*-uninfected embryos were collected, fixed, and stained with NucBlue (Hoechst 33342) for DNA, and antibodies against α- and γ-tubulin. In each panel (a column of images), channels are shown in grey scale alongside the colorized composite. Scale bars represent 5 μm. **(A)** Embryo in early prophase with one associated MTOC. **(B)** Embryo in late prophase with two associated MTOCs. **(C)** Embryo in prometaphase with five paired chromosomes (n=10). **(D)** Embryo in metaphase. **(E)** Embryo in anaphase A. **(F)** Embryo in anaphase B. **(G)** Embryo in late telophase. Additional non-pole forming MTOCs are denoted with arrowheads (C,F).

### Meiotic spindles are anastral and consistent between *Wolbachia-*infected and -uninfected embryos

To understand how *Wolbachia* induces asexual reproduction, we used immunofluorescence staining to track the events of early embryogenesis in unfertilized embryos with and without *Wolbachia* in a controlled genetic background. This builds on previous work based only on DNA staining that concluded meiosis was not impacted by *Wolbachia* (10). We considered the possibility that other defects might manifest during meiosis, prior to the chromosome segregation defect during the first mitosis, and addressed this by visualizing spindle dynamics during meiosis.

When embryos are deposited into the host, they are stalled in metaphase of meiosis I with a compact, anastral, barrel shaped spindle (Fig. 2A,D). At this point, the spindle in both *Wolbachia-*uninfected and -infected embryos had a diffuse γ-tubulin signal in a cap-like formation at the poles, consistent with acentrosomal divisions in other insects (29). Additionally, γ-tubulin was dispersed as cytoplasmic granules that did not co-localize with α-tubulin (Fig. 2A,D). Both γ-tubulin granules and polar localization were absent by meiosis II (Fig. 2B,E). During anaphase A of the division between the oocyte pronucleus and the second polar body (meiosis II), the first polar body was in prometaphase (Fig. 2B). At this point, we observed the pronuclear and polar body 2 spindle perpendicular to the oocyte surface, such that the pronuclear chromosomes were pulled toward the center of the oocyte, and polar body 2 was pulled to the periphery toward the first polar body stalled in metaphase. Diffuse γ-tubulin was also present associated with polar body 1 during Anaphase (Fig. 2B,E). After meiosis II, the pronucleus entered interphase with the haploid set of 5 chromosomes at the center of the embryo, and three compacted polar bodies at the cortex (15 chromosomes) (Fig. 2C,F). While representative images of the *Wolbachia*-uninfected and -infected embryos in interphase of NC1 varied in whether the polar bodies were associated with γ-tubulin (Fig. 2C,F), these embryos were fixed in early and late interphase, respectively, which limits direct comparison. Beyond the presence of *Wolbachia*, there were no obvious differences in meiosis between *Wolbachia*-infected and -uninfected embryos. These results support that (i) meiosis is anastral in these animals, and (ii) *Wolbachia*-mediated chromosome segregation abnormalities occur in the subsequent mitoses.

### *Trichogramma* embryos form MTOCs *de novo* during late meiosis

An important barrier to asexual reproduction in metazoans is that typically, female meiosis is acentrosomal and paternally derived centriolar MTOCs donated by the sperm are required to organize the first mitotic spindle (30, 31). A distinguishing feature of many asexual lineages is that embryos undergo a process to construct MTOCs *de novo*, which enables spindle formation during the first mitosis in the absence of fertilization (28, 32–35). However, it was unknown if *Trichogramma* shared this developmental feature. We observed that indeed, towards the end of meiosis the *Trichogramma* embryos formed cytoplasmic asters of α-tubulin, especially at the anterior of the embryo (Fig. 2G). α-tubulin asters appeared to be smallest at the most anterior end of the embryo, and larger when closer to the central nucleus. *Wolbachia*-infected embryos also formed α-tubulin asters in the same fashion (Fig. 2H). Importantly, in both the *Wolbachia*-infected and -uninfected embryos, the α-tubulin asters had clear central foci of γ-tubulin (pericentriolar material; PCM) indicating that they could function as MTOCs (Fig. 2G,H).

### Bipolar chromosome segregation occurs in the absence of *Wolbachia*

We next imaged stages of the first zygotic mitosis in *Wolbachia*-uninfected embryos. Early prophase of mitosis I in unfertilized *Wolbachia-*uninfected embryos was characterized by the recruitment of one MTOC to the nucleus (Fig. 3A). By late prophase, the nucleus had two associated MTOCs, localized to opposite poles (Fig. 3B). During prometaphase, chromosomes began to migrate to the equator, and the mitotic spindle started to form (Fig. 3C). Interestingly, in many of the embryos at this stage, each spindle pole had a clear doublet of γ-tubulin foci within the MTOC (Fig. 3C). Furthermore, some of the embryos had additional MTOCs associated with the spindle equator that were non-pole-forming (Fig. 3C). By metaphase, chromosomes were aligned at the spindle equator, and we no longer observed any γ-tubulin doublets at the spindle poles (Fig. 3D). Anaphase A featured sister chromatids in the initial stages of segregation (Fig. 3E). During anaphase, we were often able to resolve distinct γ-tubulin doublets at each spindle pole, with increased distance between them as compared to prometaphase and metaphase. By anaphase B, a distinct spindle midzone was present between partially decondensed chromosomes (Fig. 3F). By telophase, the chromosomes appeared fully decondensed, and MTOCs were associated with a reduced spindle (Fig. 3G).

During anaphase and telophase, the spindle poles almost always contained γ-tubulin doublets. In these *Wolbachia*-uninfected embryos, supernumerary MTOCs were present in 9.6% of embryos (5/52) and were typically located at the metaphase plate or spindle midzone, depending on stage (Fig. 3C,F; Supplemental Figure S3). Ultimately, mitosis I in the *Wolbachia*-uninfected embryos resulted in a successful bipolar division.

### *Wolbachia*-infected embryos form multipolar spindles

We next characterized the first mitotic division in *Wolbachia-*infected, unfertilized embryos. Similar to the *Wolbachia*-free embryos, cytoplasmic MTOCs were recruited to the nucleus during prophase (Fig. 4A). However, the prophase spindles of *Wolbachia-*infected embryos occasionally had 3 or 4 MTOCs (4/7). By late prometaphase, the condensed chromosomes were near the metaphase plate, and sister chromatid arms appeared engaged but resolved. (Fig. 4B), which aligns with *Wolbachia*-uninfected embryos at the same timepoint (Fig. 3C). In the *Wolbachia*-infected embryos that appeared to be in metaphase (as per chromosome configuration compared to *Wolbachia-*uninfected embryos, e.g., Fig. 3D), we observed two distinct categories of spindle morphology (Fig. 4C,D). Some embryos had elongated bipolar spindles, containing one MTOC at each pole (Fig. 4C). Four out of ten (40%) of the bipolar embryonic spindles in metaphase contained additional MTOCs at the equator. Alternatively, 7/15 (46.7%) of embryos with chromosomes in a metaphase configuration had multipolar spindles with one γ-tubulin focus at each pole (Fig. 4D). Of these, we observed both tripolar (n=4) and quadripolar (n=3) spindles associated with metaphase configuration chromosomes. Embryos with multipolar spindles then appeared to attempt anaphase. Within the multipolar spindle, the chromatids appeared to partially disjoin, but they remained at the center of the spindle apparatus (Fig. 4E). Following segregation failure, the chromosomes decondensed while the spindle depolymerized, as though the nucleus had transitioned to telophase (Fig. 4F). The MTOCs stayed dispersed around the nucleus (Fig. 4F).

**Figure 4.**
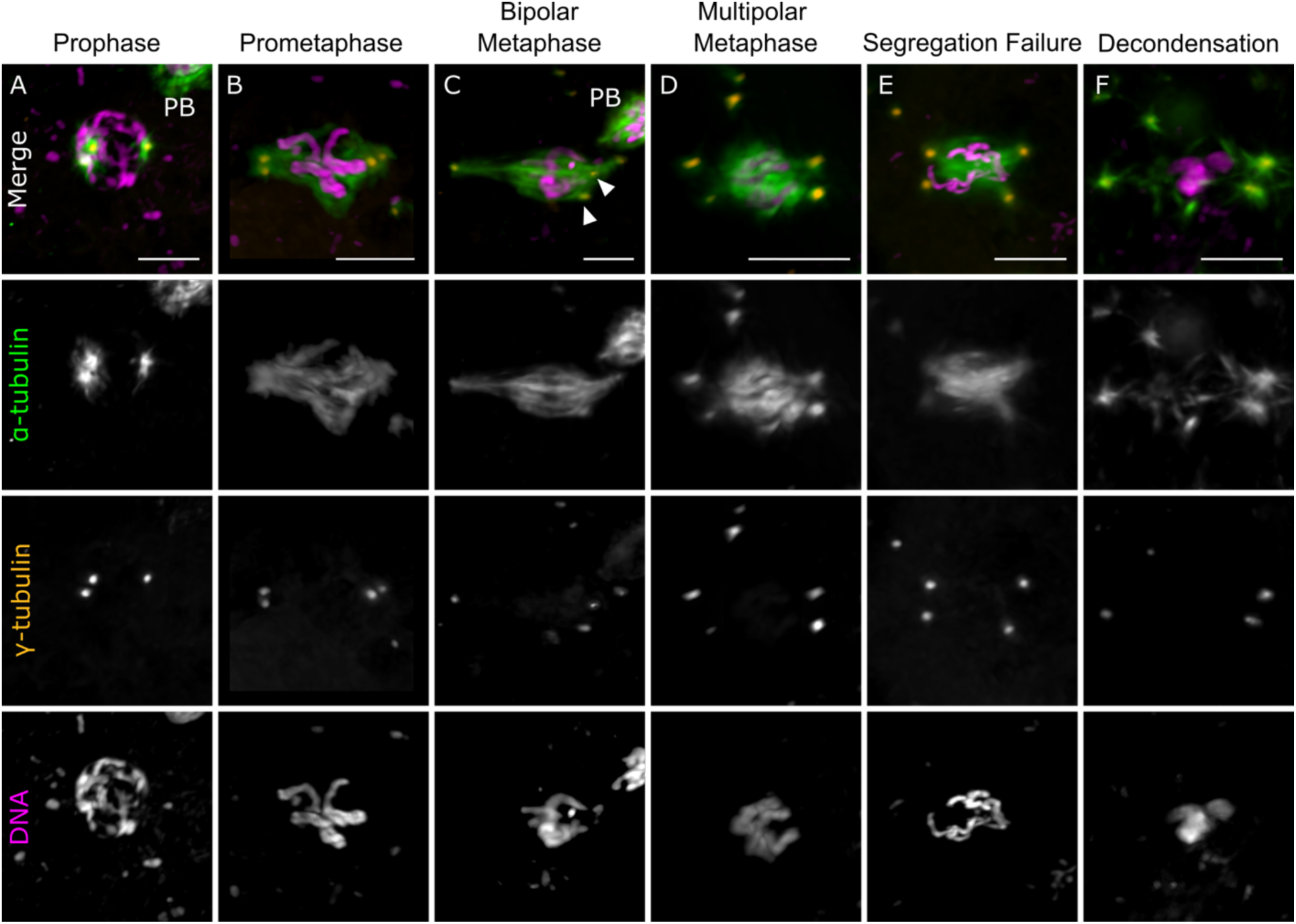
*Wolbachia*-infected embryos become multipolar during mitosis. **I.** 30- to 90-minute-old embryos were collected, fixed, and stained with NucBlue (Hoechst 33342) for DNA, and antibodies against α- and γ-tubulin. In each panel (a column of images), channels are shown in grey scale alongside the colorized composite. Scale bars represent 5 μm. Polar bodies are labeled with ‘PB’ in the composite image. **(A)** Embryo in prophase, here with three MTOCs. **(B)** Embryo in late prometaphase. **(C)** Embryo in bipolar metaphase. Two supernumerary MTOCs near the spindle equator are indicated with arrow heads. **(D)** An embryo in multipolar metaphase. **(E)** Embryo during chromosome segregation failure with a multipolar spindle. **(F)** Embryo post-mitotic failure where the nucleus decondenses.

### *Wolbachia* infection distorts the arrangement of γ-tubulin foci within the spindle

Given the combination of dynamic γ-tubulin foci (e.g., the variable presence of γ-tubulin doublets within an MTOC), and the additional poles that formed in *Wolbachia*-infected embryos, we quantified the distances between γ-tubulin foci sharing a mitotic division plane to better evaluate *Wolbachia’s* impacts on spindle morphology (Fig. 5). During prometaphase, intraplanar γ-tubulin distances were not significantly different between the *Wolbachia-*infected and *Wolbachia-*uninfected embryos (Fig. 5C; *p_adj_* = 1). At this stage, distances ranged from 0 (i.e., no distinguishable γ-tubulin doublet) to approximately 2 µm. In metaphase, all *Wolbachia-*uninfected embryos had intraplanar γ-tubulin distances of 0 µm, as no doublets were distinguishable (Fig. 5A,C). In contrast, *Wolbachia*-infected embryos in metaphase had significantly larger (Fig. 5C; *p_adj_* = 0.0381) intraplanar γ-tubulin distances (up to 7.14 µm), which reflected the diversity of the metaphase-like states, wherein both bipolar and multipolar spindle architectures were observed (Fig. 5B,C). During anaphase and telophase, the distance between γ-tubulin foci within a division plane in *Wolbachia*-infected embryos was significantly greater than in uninfected embryos (Fig. 5C; *p_adj_* = 0.0381). Importantly, we also observed that during NC1, a higher proportion of the *Wolbachia*-infected embryos were in metaphase, as compared to the *Wolbachia*-uninfected embryos (Supplemental Figure S4). This pattern could be due to a cell cycle delay, and/or this could be reflective of the significant challenge of unambiguously staging the *Wolbachia*-infected embryos given their unusual features.

**Figure 5.**
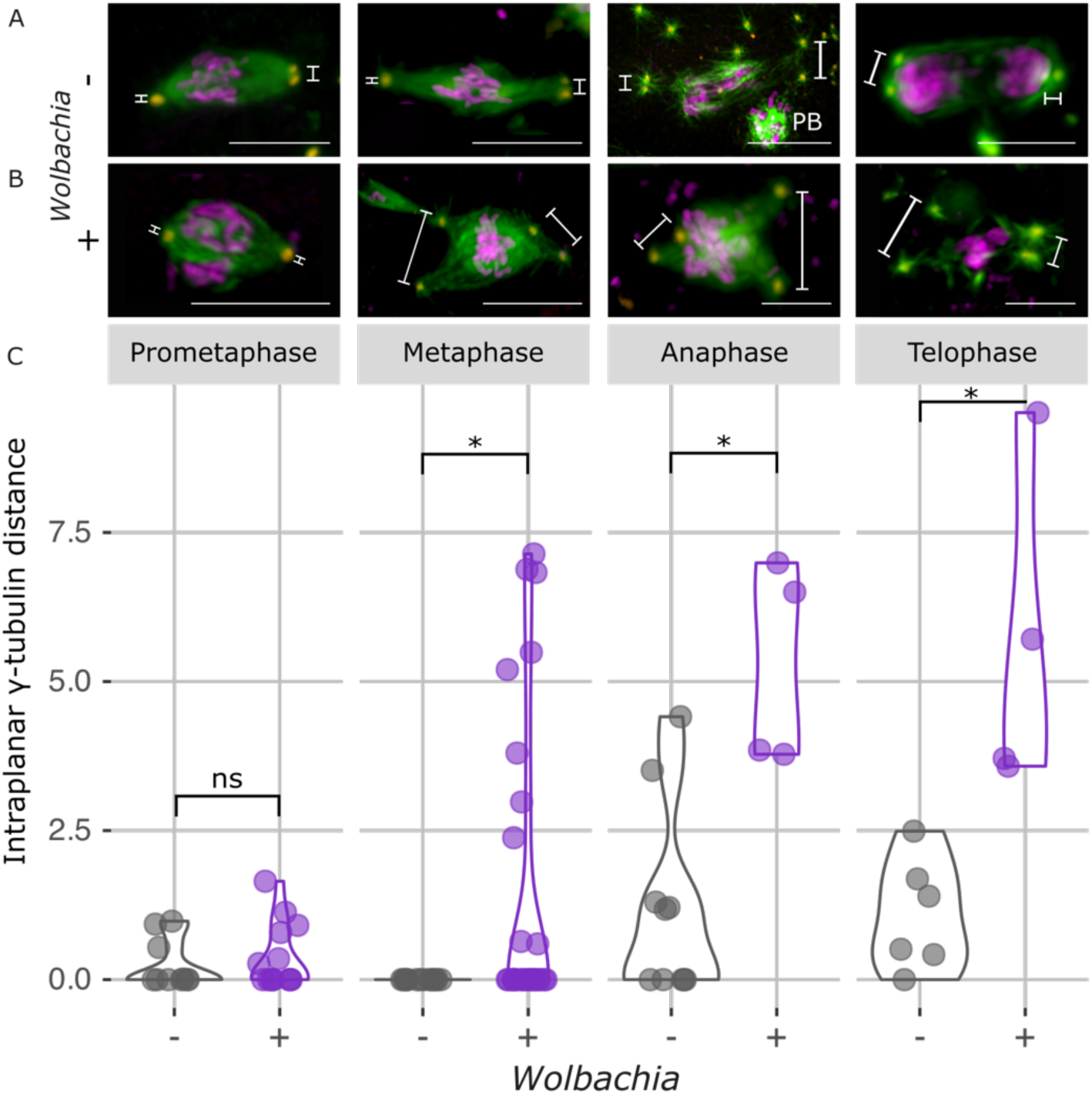
Intraplanar γ-tubulin distances increase in the presence of *Wolbachia* during the first mitosis. 30- to 90-minute-old embryos were collected, fixed, and stained with NucBlue (Hoechst 33342) for DNA, and antibodies against α- and γ-tubulin. Each image is a colorized composite. Scale bars represent 5 μm. Polar bodies are labeled with ‘PB’ in relevant images. (A, B) Representative images of mitosis I, indicating how intraplanar γ**-**tubulin distances were measured in (A) *Wolbachia*-uninfected embryos and (B) *Wolbachia*-infected embryos. (C) We measured distances between γ-tubulin foci within a division plane for *Wolbachia*-uninfected and -infected samples in prometaphase, metaphase, anaphase, and telophase of mitosis I. Staging relied on chromosome configuration as compared to the *Wolbachia*-uninfected embryos.

### Restitution mitosis spindles have supernumerary MTOCs but functionally bipolar divisions

A unique feature of *Wolbachia*-induced mitotic failure is that the animal is able to carry out subsequent, successful divisions. To understand this process, we imaged the following “restitution” mitotic cycle, NC1R. After the initial mitotic failure, chromosomes and MTOCs that would normally have segregated into two daughter nuclei were retained in the single restitution nucleus (Fig. 6A). The diploid nucleus then entered interphase and chromosomes replicated (Fig. 6B). During interphase of NC1R, restitution nuclei were associated with as many as seven MTOCs (Fig. 6B). During prophase of NC1R, the nuclei maintained associations with these numerous MTOCs, and the spindles lacked defined bipolarity (Fig. 6C). Despite the presence of supernumerary MTOCs, the prometaphase NC1R spindles that formed subsequently appeared functionally bipolar (Fig. 6D). During metaphase of mitosis NC1R, we noted the formation of pseudo-bipolar spindles (Fig. 6E). While supernumerary MTOCs were associated with the metaphase plate, they did not form strong spindle poles like in NC1 (e.g., Fig. 4E). Additionally, during metaphase of NC1R the chromosomes were neatly aligned at the equator and sister chromatid arms appeared disengaged (Fig. 6E). During anaphase A, the pseudo-bipolar spindles contained numerous MTOCs, but sister chromatids began to segregate to opposite poles (Fig. 6F). Critically, while all spindles during metaphase and anaphase of NC1R had supernumerary MTOCs, they had a dominant bipolar directionality (Fig. 6E,F). This is in contrast to spindles during the chromosome segregation failure of mitosis I that formed equal multipolar connections (Fig. 4D,E).

**Figure 6:**
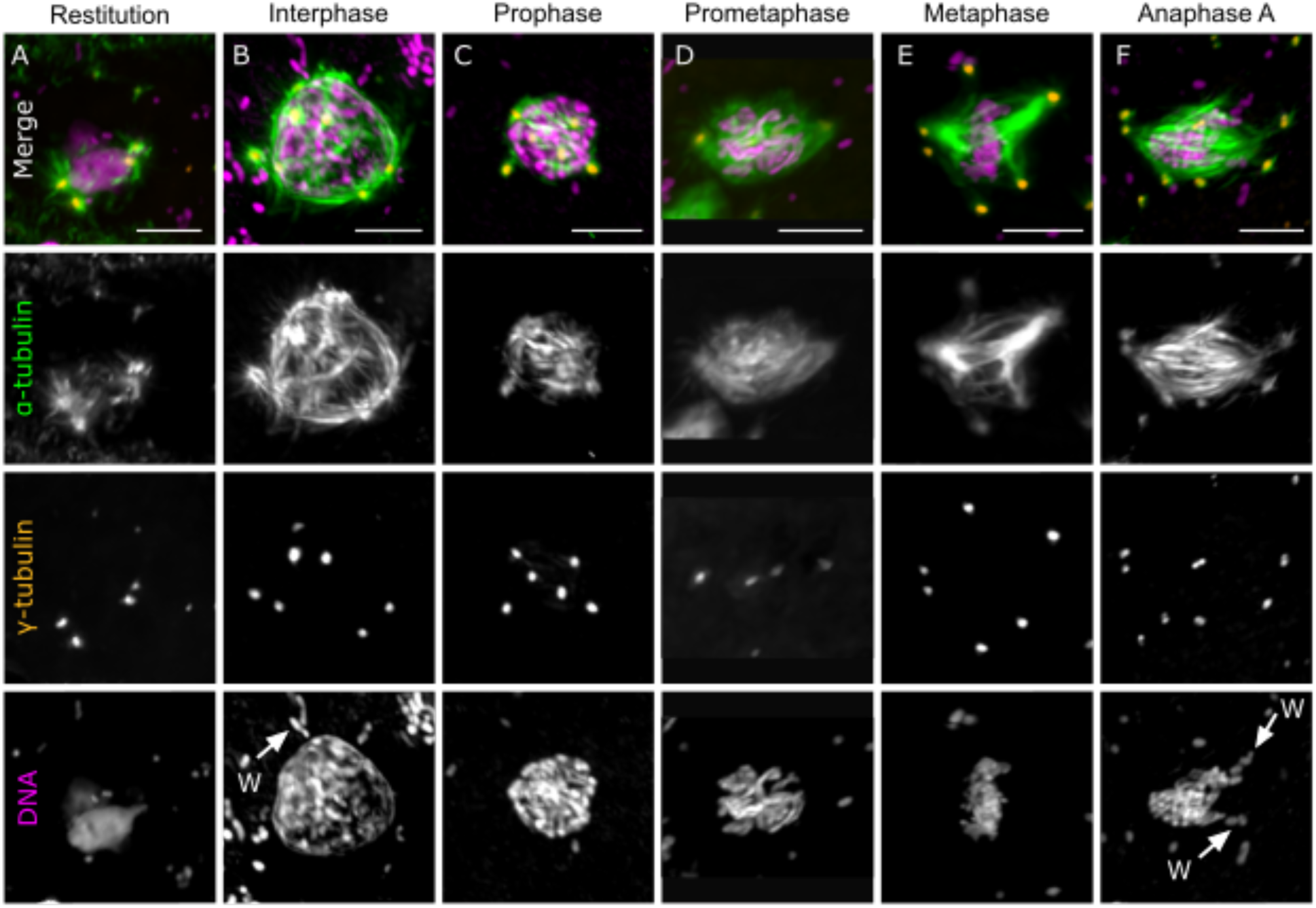
*Wolbachia-*infected embryos during the restitution mitosis have a pseudo-bipolar spindle with supernumerary MTOCs. 1- to 2.5-hour-old embryos were collected, fixed, and stained with NucBlue (Hoechst 33342) for DNA, and antibodies against α- and γ-tubulin. For each panel (a column of images), individual channels are shown in grey scale alongside the colorized composite. Scale bars represent 5 μm. *Wolbachia* that appear near the nucleus in the maximum intensity projection are labeled with “W” arrows. **(A)** Restitution nucleus of the embryo post-failure of mitosis I. **(B)** Embryo in interphase. **(C)** Embryo in prophase. **(D)** Embryo in prometaphase. **(E)** Embryo in metaphase. **(F)** Embryo in anaphase A.

### Supernumerary MTOCs persist in later mitotic divisions in *Wolbachia*-infected embryos

To understand if the supernumerary MTOCs and pseudo-bipolar spindle were persistent features of the *Wolbachia*-infected embryos, we imaged additional nuclear cycles (Fig. 7A-E) and quantified the numbers of MTOCs associated with each spindle (Fig. 7F). Here, clearly paired γ-tubulin foci that shared a single aster within one spindle pole were defined as a single MTOC. We separately analyzed the numbers of unique γ-tubulin foci associated with each spindle, regardless of their pairing (Supplemental Figure S5). During NC1, the spindles of *Wolbachia*-infected embryos were associated with 1.35x more MTOCs than the spindles of the *Wolbachia*-uninfected embryos (Fig. 7F; β = 0.33112, SE = ±0.153, z = 1.98, p = 0.048), which agrees with the presence of the multipolar spindles that occurred as early as prometaphase (Fig. 4). Specifically, 64% (27/42) of the mitotic spindles during NC1 of the *Wolbachia*-infected embryos were associated with three, four, or five MTOCs (Fig. 7C,F). In contrast, during NC1 of the *Wolbachia*-uninfected embryos, only 20% (6/30) of the spindles had aberrant numbers of MTOCs, and the additional MTOC(s) were always localized to the spindle equator (Fig, 7F, Supplemental Figure S3).

**Figure 7:**
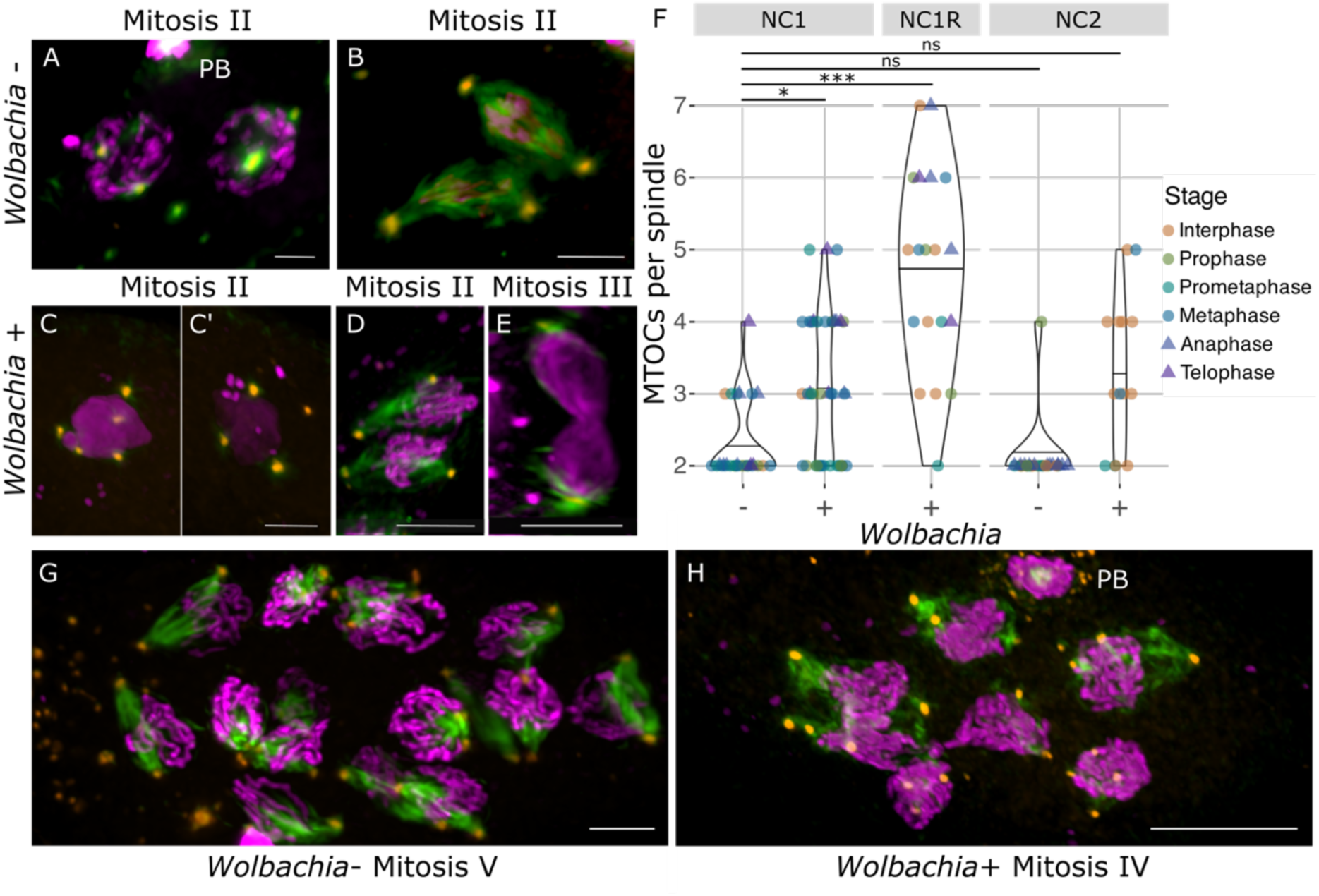
Mitotic divisions proceed despite MTOC irregularities. 1.5- to 4-hour-old embryos were collected, fixed, and stained with NucBlue (Hoechst 33342) for DNA, and antibodies against α- and γ-tubulin. Each image is a colorized composite. Scale bars represent 5 μm unless otherwise noted. **(A)** *Wolbachia*-uninfected embryo in prophase of mitosis II. Polar bodies are denoted by PB. **(B)** *Wolbachia*-uninfected embryo in Anaphase of mitosis II. **(C, C’)** *Wolbachia*-infected embryo during interphase of mitosis II. Two nuclei from the same embryo are displayed in separate panels. **(D)** *Wolbachia*-infected embryo with two dividing nuclei in prometaphase of mitosis II. **(E)** *Wolbachia*-infected embryo during telophase of mitosis III. **(F)** We quantified spindle-associated MTOCs at each stage of the cell cycle in *Wolbachia*-infected and -uninfected embryos in NC1, NC1R, and N2. Clearly paired γ-tubulin foci were classified as a single MTOC based on proximity and sharing an aster. **(G)** *Wolbachia*-uninfected embryo in prometaphase of mitosis V. The scale bar represents 10 μm. **(H)** *Wolbachia*-infected embryo during prometaphase of mitosis IV. Polar bodies are labeled with PB. The scale bar represents 10 μm.

In the next NC (NC2 for *Wolbachia*-uninfected embryos, NC1R for *Wolbachia*-infected embryos), the numbers of spindle associated MTOCs depended on *Wolbachia*-infection. In NC2, almost all *Wolbachia*-uninfected embryos (31/32) had two MTOCs associated with each spindle (Fig. 7A,B,F). Conversely, during NC1R in the *Wolbachia*-infected embryos, the number of spindle-associated MTOCs significantly increased (2.08x) relative to the NC1 (β = 0.764, SE = ±0.164, z = 4.52, p < 0.0001). This likely reflects the inheritance of MTOCs from the failed NC1 and subsequent MTOC replication. Interestingly, in NC2, while some spindles of *Wolbachia*-infected embryos still had supernumerary MTOCs, the numbers of MTOCs per spindle were only marginally significantly different as compared to NC1 (β = 0.398, SE = ±0.193, z = 2.16, p = 0.056). Overall, the presence of *Wolbachia* was associated with an increased frequency of supernumerary MTOC associations in NC1 and NC1R. This pattern was consistent with the numbers of γ-tubulin foci associated with each spindle, regardless of whether they were paired within an MTOC (Supplemental Figure S5).

Finally, we imaged even later NC (up to mitosis V) and saw that *Wolbachia*-uninfected embryos still had the expected two MTOCs per nucleus and formed bipolar spindles (Fig. 7H). Additionally, we noticed that the nuclei of *Wolbachia*-uninfected embryos divided synchronously; for example, all 16 nuclei in the representative image of the fifth mitotic cycle are in prometaphase (Fig. 7H). In contrast, the spindles of *Wolbachia*-infected embryos were frequently associated with supernumerary MTOCs, and in many cases daughter nuclei were associated with different numbers of MTOCs (Fig. 7C). Furthermore, the divisions in *Wolbachia*-infected embryos appeared slightly asynchronous, as evidenced by varying levels of spindle formation during mitosis IV (Fig. 7G). Despite these irregularities, bipolar spindles formed, and divisions were successful (Fig. 7D-G). Curiously, the cytoplasmic MTOCs persisted across all later stages that we observed, though only rarely did they form spindle connections in the absence of *Wolbachia.* Ultimately, across subsequent cycles, the mitotic divisions of the *Wolbachia*-infected embryos started to converge on a much more organized configuration akin to the *Wolbachia*-uninfected embryos.

## DISCUSSION

The discovery that *Wolbachia* can induce the asexual production of female offspring (9), via selective impairment of the first mitotic division of unfertilized *Trichogramma* embryos (10) was made more than three decades ago. However, the underlying mechanism has since remained a mystery, despite many other reports of *Wolbachia-*mediated gamete duplication in other species (15, 36). As such, we examined differences between *Wolbachia*-infected and -uninfected embryos to uncover the structural basis of mitotic failure. In the absence of *Wolbachia*, meiosis and early embryonic mitoses in *Trichogramma* share many similarities with other animal models including mammals, *Drosophila,* and other parasitoid wasps (29, 30, 37). The meiotic divisions are anastral and give rise to three polar bodies and the oocyte pronucleus. As expected, meiosis in the *Wolbachia*-infected embryos closely mirrored meiosis in the *Wolbachia*-uninfected embryos (10). While γ-tubulin is associated with the first meiotic division, it does not form distinct foci as would be found in astral spindles (38). In all embryos, cytoplasmic asters of microtubules with γ-tubulin foci form towards the end of meiosis through a process known as *de novo* MTOC biogenesis (34). At least two of these *de novo* MTOCs associate with the nucleus during prophase to facilitate the first mitotic division (39). Embryonic mitoses in *Trichogramma* have features characteristic of rapid syncytial divisions including the separation of paired γ-tubulin foci (indicative of centriolar disengagement; (40)), and the early decondensation of chromosomes, both of which facilitate more rapid entry into the next nuclear cycle (41). However, in the *Wolbachia*-infected embryos we characterized three differences during the first embryonic mitosis: (1) increased instances of supernumerary MTOCs, (2) a transition to multipolar spindles, and (3) an incomplete segregation of chromosomes.

First, we observed that *Wolbachia-*infected embryos in NC1 were more likely to have supernumerary MTOCs. Despite this, a bipolar mitotic spindle initially forms and chromosomes appear to be bioriented, though multipolar spindles appear as early as prometaphase. In metaphase, the chromosomes are morphologically comparable to those of the *Wolbachia*-uninfected embryos at this stage. A number of *Wolbachia*-infected embryos had what appeared to be a typical, bipolar spindle, suggesting they do have a “true” metaphase for a period of time. We also observed metaphase-stage chromosomes with tripolar and quadripolar spindles in the *Wolbachia-*infected embryos, suggesting that spindle defects largely manifest during metaphase. Additionally, a greater proportion of *Wolbachia*-infected embryos in mitosis I were in metaphase as compared to uninfected embryos, suggesting an increased metaphase duration. During chromosome segregation failure, the MTOCs within a division plane further increase in separation as chromatids appear to partially disjoin but fail to segregate to opposite poles, likely due to the absence of appropriate spindle tension (42, 43). Notably, we never observed a distinct anaphase spindle in *Wolbachia*-infected embryos during the first mitosis, and all spindles at this later stage appeared to be quadripolar. The chromosomes bypassed segregation and decondensed as they would during telophase, but then reformed into a single restitution nucleus, characteristic of mitotic slippage (Fig. 8) (42). We have two alternative hypotheses for the origin of multipolar spindles: (1) over-recruitment of MTOCs, and (2) MTOC splitting.

**Figure 8:**
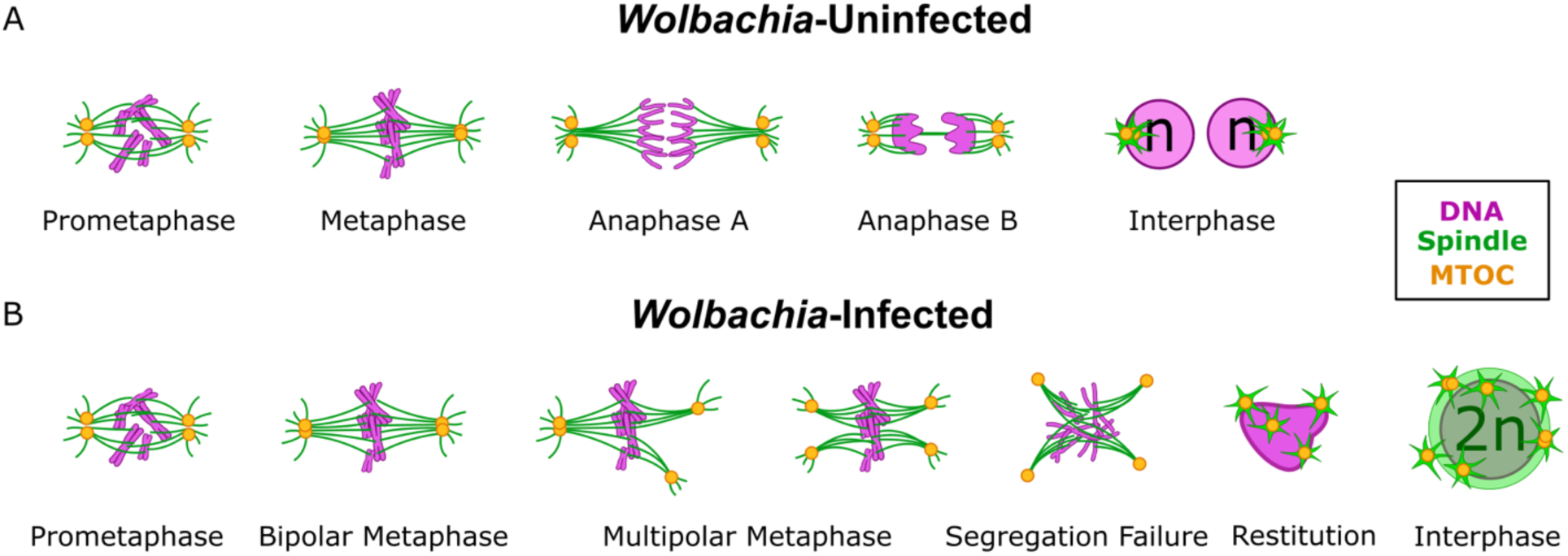
Mechanistic model of *Wolbachia*-mediated mitotic failure. **(A)** Schematic of mitotic progression through NC1 into NC2 of unfertilized, *Wolbachia*-uninfected embryos. The prometaphase spindle is bipolar, and chromosomes migrate to the spindle equator, while a slight distance between γ-tubulin foci within each polar MTOC is distinguishable. When metaphase chromosomes align at the spindle equator, paired γ-tubulin foci within each MTOC are indistinct. During anaphase A, sister chromatids segregate as distances between γ-tubulin foci within each pole increase. Chromosomes during Anaphase B separate further and begin to decondense while a spindle midzone forms. Nuclei in the interphase of mitosis 2 are haploid and each receive a pair of γ-tubulin foci. **(B)** Schematic of mitotic progression during early development of unfertilized, *Wolbachia*-infected embryos. The bipolar spindle of NC1 prometaphase pulls chromosomes to the equator, with slight distance between γ-tubulin foci within each polar MTOC. The NC1 metaphase spindle is initially bipolar and aligns chromosomes at the metaphase plate. While chromosomes remain in a metaphase conformation, the additional spindle poles form. During segregation failure, chromosomes remain near the center of the mitotic apparatus and partially disjoin. At this point, the NC1 spindle is made up of 3-4 poles, each containing an unpaired MTOC. The spindle depolymerizes, and DNA decondenses to form a mass with associated MTOCs. The now diploid nucleus reorganizes, enters NC1R, undergoes DNA replication, and MTOC numbers increase.

The significant increase in the number of spindle-associated MTOCs in the *Wolbachia-*infected embryos as early as prophase of NC1 suggests that MTOC over-recruitment might be responsible for the formation of the multipolar spindles. Importantly, supernumerary MTOCs seem to be a feature of *Trichogramma* development more generally, as they also occurred in NC1 of the *Wolbachia*-uninfected embryos. In other systems, supernumerary MTOCs are often clustered at spindle poles (44), but in *Wolbachia*-uninfected *Trichogramma* they seem to remain at the spindle equator and do not form a pole. *Trichogramma* may have mechanisms for not only regulating the recruitment of MTOCs but also licensing an appropriate number of associated MTOCs to form spindle poles, and *Wolbachia* might target either of these processes. However, many of the *Wolbachia*-infected embryos reached metaphase with only two MTOCs, and we never observed canonical anaphase. So, if MTOC over-recruitment is responsible, this would necessitate a very rapid recruitment of additional MTOCs in a relatively short window of time. However, given that the recruitment of *de novo* MTOCs is a unique feature of the first mitotic division of the embryo, this could explain how *Wolbachia*-mediated mitotic failure only manifests in the first division, and only in unfertilized embryos that do not receive canonical centrioles from the sperm.

Alternatively, MTOC recruitment and the initial formation of a bipolar spindle may not be the target of *Wolbachia*. Instead, defects impacting the two already pole-forming MTOCs might lead to the formation of a multipolar spindle. For example, polar MTOCs might undergo an inappropriate duplication and/or premature disengagement. Several observations support the hypothesis of premature disengagement. First, we never observed more than four spindle poles in the *Wolbachia*-infected embryos. Second, pole-forming MTOCs in the *Wolbachia*-uninfected embryos contained paired γ-tubulin foci. However, in the *Wolbachia*-infected embryos, if a mitotic division plane had two spindle poles, the MTOCs associated with each of those poles only ever had a single γ-tubulin focus, reflecting a loss of MTOC integrity. Importantly, in the *Wolbachia*-uninfected embryos, the paired γ-tubulin foci within a polar MTOC do separate during anaphase, consistent with normal centriolar disengagement (40). In the *Wolbachia*-uninfected embryos the γ-tubulin foci separate to a maximum of 4.3 µm during anaphase and telophase, and they are still associated with a single spindle pole. In contrast, in the *Wolbachia*-infected embryos the minimum distance between γ-tubulin foci within the same plane (i.e., different spindle poles on the same side of the chromosomes), was just under 4 µm, and maximum distances approached 10 µm: well beyond the level of γ-tubulin foci separation seen in *Wolbachia*-uninfected controls.

Importantly, while premature disengagement could explain the formation of multiple spindle poles, it is not clear if this is the proximate cause of chromosome segregation failure, or, a symptom of some other *Wolbachia*-mediated effect. For example, the primary defect might be direct interference with chromosome disjunction, and MTOC disengagement and/or multipolarity could be by-products. Under standard models of mitosis, once chromosomes are aligned at the metaphase plate with bipolar kinetochore attachment, the spindle assembly checkpoint would be satisfied, and the anaphase promoting complex/cyclosome (APC/C) is activated (45). The APC/C then triggers the degradation of securin, releasing separase that takes part in both sister chromatid disjunction and centriole disengagement by the cleavage of cohesin (45). *Wolbachia* may target regulators of chromatid disjunction, inhibiting the cleavage of cohesin and leaving the mother and daughter MTOC pair to prematurely disengage through separase-independent mechanisms, taking advantage of a less sensitive spindle assembly checkpoint characteristic of syncytial divisions (46, 47). Our observations of inconsistent chromatid disjunction and MTOC disengagement during mitotic failure could both be explained by cohesion fatigue following mitotic delay. However, a model in which impaired chromatid disjunction is the primary driver of the *Wolbachia*-mediated mitotic failure is more challenging to reconcile with the fact that the failure is restricted to the first embryonic mitosis, and only in unfertilized embryos (48, 49).

In contrast, *Wolbachia* might act directly on MTOCs or the associated regulatory processes, and failed chromatid segregation could be the by-product. We hypothesize that there are unique features of the *de novo* MTOCs and their maturation that could enable *Wolbachia* to uniquely target the first embryonic mitosis and prevent *Wolbachia* from acting on canonical sperm-derived MTOCs. However, very little is known about the process of *de novo* MTOC biogenesis generally, let alone in *Trichogramma*. How these structures are assembled, how or when they undergo replication, and how recruitment of an appropriate number of MTOCs is regulated are all unknown. In distantly related hymenopterans, *de novo* MTOCs are associated with at least one canonical centrosomal protein (CP190) (34), but whether this is conserved in *Trichogramma*, and whether the *de novo* MTOCs indeed contain centrioles (and how many) is unknown. Under canonical centrosomal maturation and assembly, the mother and daughter centriole are held together by a proteinaceous linker within a PCM formed by a matrix of proteins with microtubule-nucleating and -anchoring capabilities (50). While we are able to track PCM via γ-tubulin, our experiments cannot determine if all γ-tubulin foci contain canonical centrosomal and/or centriolar proteins. Given that *Trichogramma* are quite diverged from model systems for which numerous antibodies are available and reverse genetics is possible, significant tool development will be required to address questions focused on the mechanisms of *de novo* MTOC biogenesis in this system.

Curiously, mitosis of NC1R is the first successful division of the embryo but is morphologically distinct from all mitotic divisions of the *Wolbachia*-uninfected embryos, and from the first mitosis of the *Wolbachia*-infected embryos. Specifically, while we saw that mitosis of NC1R almost always had supernumerary MTOCs, the spindle that formed was effectively bipolar. In other animals there are mechanisms for clustering extra centrosomes that allow the cell to maintain a pseudo-bipolar mitotic spindle and undergo chromosome segregation (44). In the *Wolbachia*-infected embryos, there appears to be some level of clustering of supernumerary MTOCs at the spindle poles during anaphase of NC1R, wherein microtubules bundle to a focal point at each pole and multiple MTOCs extend from that focal point. We also noticed that the numbers of spindle-associated MTOCs during NC1R did not align with what we would expect if MTOCs were strictly inherited from NC1. For example, if NC1 fails with a quadripolar spindle, there should be four MTOCs in NC1R that then undergo replication during interphase. However, many spindles in NC1R and NC2 had odd numbers of MTOCs, suggesting that either (1) MTOC-spindle associations are potentially transient in nature, and/or (2) not all MTOCs undergo replication each nuclear cycle. Regardless, that *Trichogramma* embryos generally seem permissive to supernumerary MTOCs and relatively disorganized spindles may underlie their ability to recover from the *Wolbachia*-mediated mitotic failure.

While we have determined there are MTOC and spindle abnormalities due to *Wolbachia* infection, the precise molecular interactions occurring between *Wolbachia* and *Trichogramma* remain in question. We previously identified two putative parthenogenesis inducing *Wolbachia* proteins, PifA and PifB (51). PifA and/or PifB orthologs are present in a suite of PI-*Wolbachia* strains that infect *Trichogramma* and several other genera of parasitoid wasps (36, 51–54). Of note, PifB contains a large leucine rich repeat (LRR) domain that has structural homology to LRR receptor-like serine/threonine protein kinases and F-box proteins including F-box/LRR protein 18, suggesting a role of PifB in regulating mitotic events (51). Alternatively, PifA contains a nucleoporin-like domain and localizes to the poles of dividing nuclei in yeast heterologous expression assays (51). During mitosis, nucleoporins often relocalize to spindle poles or kinetochores, which suggests PifA may be capable of impacting mitosis (55). Regardless, the restriction of the failure to the first mitosis of unfertilized embryos strongly points to a *Wolbachia* effector protein that can target unique aspects of initiating sperm-independent zygotic mitosis.

In summary, we demonstrated that *Wolbachia-*mediated mitotic failure is characterized by supernumerary MTOCs, a transition to multipolar spindles, and incomplete chromosome disjunction. This research establishes a fascinating model system for understanding not only asexual reproduction and microbial impacts on the cell cycle, but also *de novo* MTOC biogenesis, and the constraints and flexibility of mitosis.

## ACKNOWLEDGEMENTS

Research reported in this publication was supported by the National Institute of General Medical Sciences of the National Institutes of Health under award number R35GM150991 to ARIL and the National Institute of Food and Agriculture Predoctoral Fellowship to LCF (NIFA 2023-67034-40496). This work was supported by the resources and staff at the University of Minnesota University Imaging Centers (UIC), RRID: SCR_020997.

