## Supplemental Table and Figures for "Multipolar spindle assembly and mitotic slippage underlie symbiont-mediated asexual reproduction"

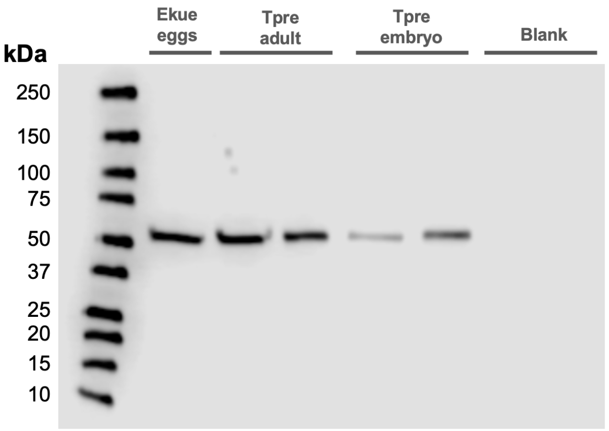


**Supplemental Figure S1**. Western blot for α-tubulin. Mass is indicated in kilodaltons. Ekue = *Ephestia kuehniella* (host eggs); Tpre = *Trichogramma pretiosum*.

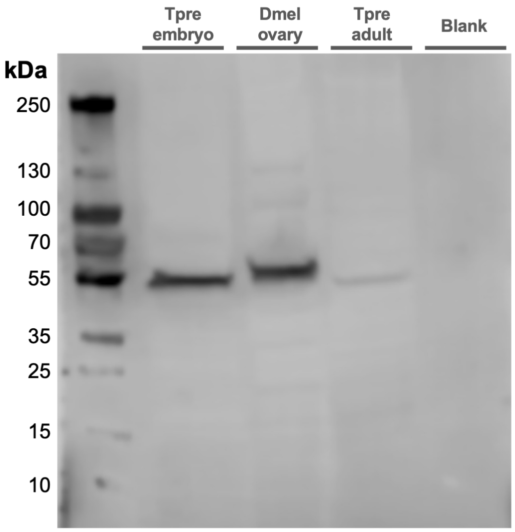


**Supplemental Figure S2**. Western blot for γ-tubulin. Mass is indicated in kilodaltons. Tpre = *Trichogramma pretiosum*; Dmel = *Drosophila melanogaster*.

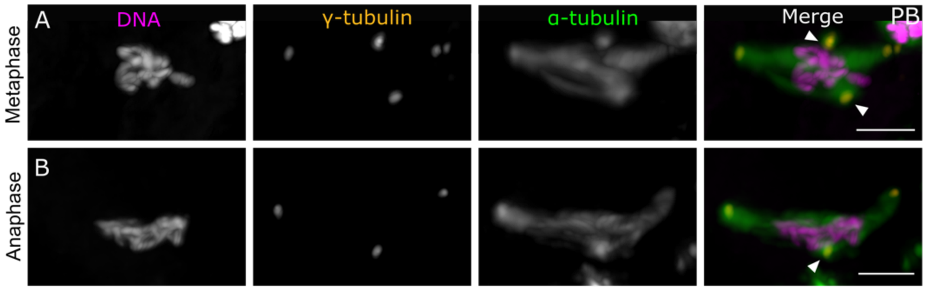


**Supplemental Figure S3. *Wolbachia*-uninfected embryos have occasional supernumerary MTOCs.** 0- to 2-hour-old embryos were collected, fixed, and stained with NucBlue (Hoechst 33342) for DNA, and antibodies against α- and γ-tubulin. In each panel, channels are shown in grey scale alongside the colorized composite. Scale bars represent 5 µm. **(A)** An embryo in metaphase, where the spindle has a two additional MTOCs at the equator. **(B)** An embryo in anaphase, where an additional MTOC is present at the spindle midzone. Arrowheads indicate the extra MTOC association.

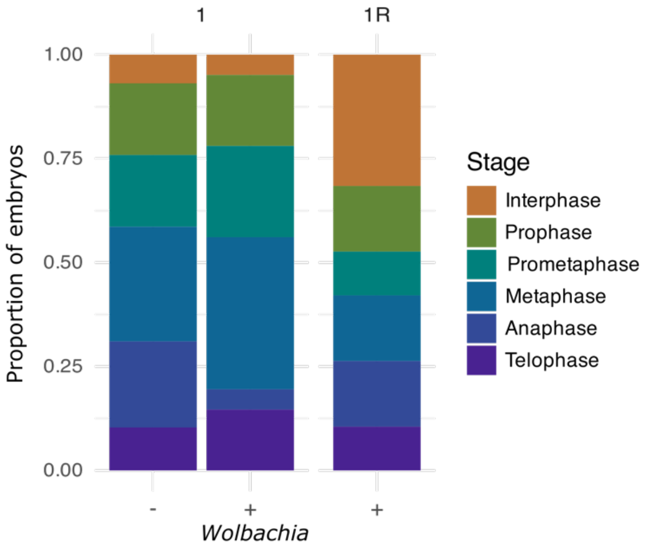


**Supplemental Figure S4. *Wolbachia*-infected embryos during NC1 have a greater proportion of nuclei in metaphase**.

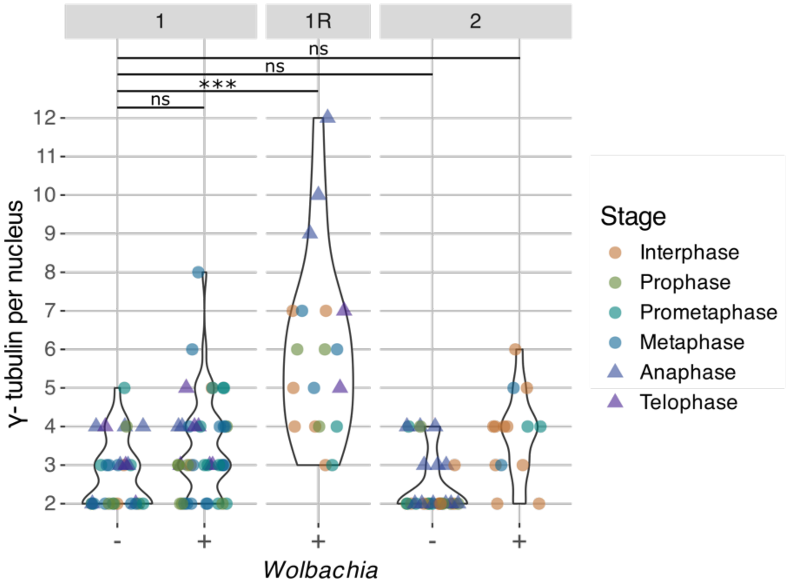


**Supplemental Figure S5: γ-tubulin foci associated with the spindle are affected by *Wolbachia* infection.** We quantified spindle-associated γ-tubulin foci at each stage of the cell cycle in *Wolbachia*-infected and -uninfected embryos in NC1, 1R, and 2. The triangles indicate embryos in anaphase and telophase, where failure occurs in mitosis I of *Wolbachia*-infected embryos.

**Supplemental Table S1: Key reagents and resources**

| REAGENT or RESOURCE | SOURCE | IDENTIFIER |
|---|---|---|
| **Antibodies** | | |
| Alpha Tubulin Monoclonal antibody IgG2b | ProteinTech | RRID:AB_11042766; Cat#66031-1; Clone#1E4C11 |
| Gamma Tubulin Polyclonal Antibody | Invitrogen | Cat#PA5-34815; RRID:AB_2552167 |
| Nano-Secondary® alpaca anti-mouse IgG2b, recombinant VHH, Alexa Fluor® 488 | ProteinTech | RRID:AB_2827581; Cat#sms2bAF488-1; Clone#VHH0275, VHH0288 |
| ChromoTek Nano-Secondary® alpaca anti-human IgG/anti-rabbit IgG, recombinant VHH, Alexa Fluor® 568 | ProteinTech | RRID:AB_2827586; Cat#srbAF568-1 Clone#CTK0101, CTK0102 |
| Goat α-Mouse IgG (H+L) Cross-Adsorbed Secondary Antibody, HRP | Life Technologies | RRID:AB_2536527 Cat#G-21040 |
| Goat α-Rabbit IgG (H+L) HRP-conjugated Affinipure | ProteinTech | RRID: AB 2722564 Cat#SA00001-2 |
| **Bacterial strains** | | |
| *Wolbachia* Endosymbiont of *Trichogramma pretiosum* | Insectary line (19) | N/A |
| **Experimental models: Organisms/lines** | | |
| *Trichogramma pretiosum* Introgressed Line B Infected (IILB+) | Generated (18) | N/A |
| *Trichogramma pretiosum* Introgressed Line B Uninfected (IILB-) | Generated (18) | N/A |
| *Drosophila melanogaster* DGRP-352 | Generated 6/11/26 12:30:00 PM | RRID:BDSC_83728 |
| *Ephestia kuehniella* | Koppert Global or Bioline Agrosciences | N/A |
| **Oligonucleotides** | | |
| W-specF: 5′-CATACCTATTCGAAGGGATA −3′ | Generated (27) | N/A |
| W-specR: 5′-AGCTTCGAGTGAAACCAATTC −3′ | Generated (27) | N/A |
| **Software** | | |
| NIS Elements | Nikon Instruments Inc. | RRID: SCR_014329 Version 6.10.01 |
| RStudio | https://www.R-project.org/ | Version 4.4.1 |
| Inkscape | https://www.inkscape.org/ | Version 1.3.2 |
